# Antidiabetic “gliptins” affect biofilm formation by *Streptococcus mutans*

**DOI:** 10.1101/157503

**Authors:** Arpan De, Arianna Pompilio, Jenifer Francis, Iain C. Sutcliffe, Gary W. Black, Giulio Lupidi, Dezemona Petrelli, Luca A. Vitali

## Abstract

*Streptococcus mutans*, a dental caries causing odontopathogen, produces X-prolyl dipeptidyl peptidase (Sm-XPDAP, encoded by *pepX*), a serine protease known to have a nutritional role. Considering the potential of proteases as therapeutic targets in pathogens, this study was primarily aimed at investigating the role of Sm-XPDAP in contributing to virulence-related traits. Dipeptidyl peptidase (DPP IV), an XPDAP analogous enzyme found in mammalian tissues, is a well known therapeutic target in Type II diabetes. Based on the hypothesis that gliptins, commonly used as anti-human-DPP IV drugs, may affect bacterial growth upon inhibition of Sm-XPDAP, we have determined their *ex vivo* antimicrobial and anti-biofilm activity towards *S. mutans*. All three DPP IV drugs tested reduced biofilm formation as determined by crystal violet staining. To link the observed biofilm inhibition to the human-DPP IV analogue present in *S. mutans* UA159, a *pepX* isogenic mutant was generated. In addition to reduced biofilm formation, CLSM studies of the biofilm formed by the *pepX* isogenic mutant showed these were comparable to those formed in the presence of saxagliptin, suggesting a probable role of this enzyme in biofilm formation by *S. mutans* UA159. The effects of both *pepX* deletion and DPP IV drugs on the proteome were studied using LC-MS/MS. Overall, this study highlights the potential of Sm-XPDAP as a novel anti-biofilm target and suggests a template molecule to synthesize lead compounds effective against this enzyme.

## Introduction

Dental caries is the most prevalent, multifactorial, globally increasing oral health problem among children and adults (Selwitz et al., 2007; Bagramian et al., 2009). It is a manifestation of biofilm formation by certain members of the indigenous oral microbiota that are aciduric and acidogenic. Among them, *Streptococcus mutans* is one of the key etiological agents of dental caries. *S. mutans* is known to code for several peptidases and exoglycosidases that can facilitate utilization of human saliva as a source of nutrition (Ajdić et al., 2002). X-prolyl dipeptidyl aminopeptidase (XPDAP) is one such narrow substrate range cytoplasmic endopeptidase found in *S. mutans*, which help in utilization of proline-rich salivary polypeptides (Cowman and Baron, 1993; Cowman and Baron, 1997). Collagenolytic and caseinolytic activities demonstrated by XPDAP further suggests its importance as a putative virulence factor in *S. mutans* (Cowman *et al.*, 1975; Rosengren & Winblad, 1976). In *Streptococcus suis* and *Streptococcus gordonii* extracellular XPDAP play a role also in cellular invasion (Ge et al., 2009; Goldstein et al., 2001). Similarly, periodontal pathogen *Porphyromonas gingivalis* also had altered virulence in absence of XPDAP (Yagishita et al., 2001).

An analogous enzyme to XPDAP, DPP IV is also found in mammalian tissues and is a target for maintaining glucose homeostasis in Type II diabetic patients (Green et al., 2006; Cowman and Baron, 1997). Diabetes, an abnormal metabolic disorder, is an epidemic of significant healthcare concern among both developed and developing countries (King et al., 1998). Certain drugs, namely saxagliptin, vildagliptin and sitagliptin, are commonly used anti-human DPP IV (AHD) molecules used in the treatment of Type II diabetes (Green et al., 2006). DPP IV targets incretin hormones such as GLP-1, thereby decreasing their plasma levels. Inhibition of DPP IV leads to the opposite effect, which results in a restoration of glucose homeostasis in diabetic patients (Wang et al., 2012). As a novel approach, our recent investigation has shown that *S. mutans* XPDAP (Sm-XPDAP, encoded by *pepX*) is inhibited by saxagliptin *in vitro* (De et al., 2016). In an extension of this work and hypothesising a probable role of Sm-XPDAP in virulence, herein we have evaluated the *ex vivo* effect of these molecules on cell growth and biofilm formation by *S. mutans*. A *pepX* (SMU.35) isogenic mutant was also generated. Furthermore, whole cell proteome analysis of AHD treated cells and the isogenic mutant was performed to identify possible consequences of Sm-XPDAP inhibition or deletion.

## Methods

### MIC and Biofilm formation assay

The MIC was determined by microdilution assay according to the Clinical and Laboratory Standards Institute (CLSI, 2011), with the exception of the medium used, which was BHI (da Silva et al., 2013; Ahn et al., 2012). The highest concentration of 2048µg/mL of each AHD was serially diluted down to 4µg/mL and the final density of mid-exponential phase cells used was 10^6^ CFU/mL. The drugs were dissolved in sterile water and erythromycin was used as a positive control.

Biofilm formation was assessed by a semi-quantitative crystal violet method in polystyrene 96-well cell culture plates (Costar 3595; Corning Inc., NY) as previously described (Ahn et al., 2008). An overnight culture of *S. mutans* was transferred into pre-warmed BHI and grown till mid-exponential phase and then diluted 50 fold in semi-defined biofilm medium (SDM) supplemented with 20mM glucose or sucrose. Aliquots (100µL) of this culture were added to serially diluted drug in water (2048µg/mL to 4µg/mL), to make a final volume of 200µL (with a 100-fold final dilution of cells) and incubated for 20 hours. The adhered cells were stained with 1% crystal violet for 15 minutes and the extract was quantified at 495nm. The viability of the cells in the culture medium (planktonic phase over the formed biofilm) at each concentration of drug was determined by CFU counting.

### Construction of *pepX* deletion mutants

A *pepX* deletion mutant was generated by PCR ligation mutagenesis method (Lau et al., 2002). Briefly, erythromycin cassette, upstream and downstream flanking regions (about 600bps) of *pepX* was amplified using specific primers (Table 2). The amplicons were digested using NcoI and SacI and ligated at 16⁰C overnight. The resulting ligation mixes were used for PCR to obtain a mutagenic construct using primers pepX-Up-F and pepX-Dn-R. This fragment was naturally transformed into mid-log phase *S.mutans* UA159, grown in Todd-Hewitt broth containing 10% sucrose and the recombinants were selected on BHI agar containing 10µg/mL erythromycin (Petersen and Scheie, 2000).*The pepX* mutant was confirmed by colony PCR and Sanger sequencing.

**Table 2.**
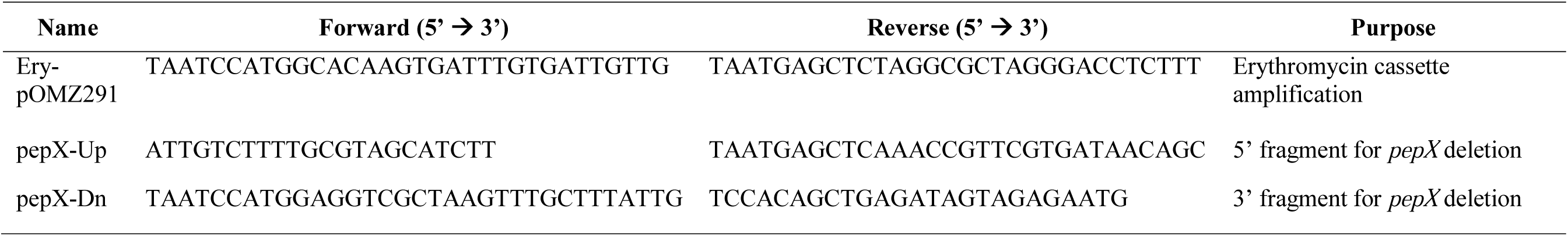
Oligonucleotide DNA primers used in the present study.

### CLSM examination of Biofilm

The structure of biofilm of the *pepX* mutant and wild-type *S. mutans*, grown on polystyrene discs in the presence of SDM and glucose and AHD drugs as described above, was evaluated using an LSM-510-META laser scanning microscope attached to an Axioplan-II microscope (Zeiss). The non-adherent cells were washed in saline, biofilms were stained with Live/Dead BacLight-(1X) (Molecular Probes Inc.) for 20 minutes and rinsed three times in saline to remove excess stain. Subsequently, the stained discs were examined with an alpha Pan-Fluor 100X objective under excitation at 488nm (Argon-laser) and 543nm (He-Ne-laser), and emission filter ranging 585-615nm and 505-530nm for Propidium iodide and SYTO9, respectively. ImageJ v1.48 (NIH, USA) was used to process images. The proportion of viable cells (green) versus dead cells (red) was determined based on the intensity at each pixel using ImageJ (Nance et al., 2013). The proportion of green signal and the red signal was calculated by multiplying the total number of pixels with the given intensity (0–255) at each channel and then dividing it by the sum of the intensity value for each signal measured at each image stack.

### Proteome analysis of biofilm-grown cells

The proteome of biofilm-grown cells in absence or presence of an AHD drug and that of the Δ*pepX* mutant were analysed from 48mL of culture. In brief, after 20 hours incubation, the harvested biofilm cells were lysed. The lysate was used to acetone precipitate all proteins overnight at −20°C. The precipitate was washed in 80% and, successively, in 40% acetone, air dried and then partially pre-fractionated by 1D PAGE. The protein containing gel was stained, destained and sectioned into pieces for treatment in 100mM NH_4_HCO_3_ and acetonitrile (ACN) for complete removal of Coomassie stain. The gel slices were dehydrated in ACN and digested in 40µg/mL trypsin (Trypsin Gold, MS Grade, Promega) at 37°C overnight. Digestion was then stopped by adding 50% ACN (v/v) and 5% formic acid (v/v) with shaking for 30 min. The peptides-containing digested extract was removed and the gel pieces were further extracted in ACN and formic acid before freeze drying. The lyophilized peptide digest was mixed in 5% ACN and 0.1% formic acid (v/v) and then run in a LC-NanoPump coupled to a tandem mass spectrometer (Thermo-Q-Exactive attached to HPLC Ultimate-3000-RSLCnano system), through an Easy Spray C18 column (PepMap-RSLC, 75µm×500mm, Thermo Scientific) in a gradient solvent mixture of water and ACN containing 0.1% formic acid. The run was carried out for 215 minutes at a flow rate of 0.3µL/min with a scan range of 350–1800 m/z. The mass spectrum data (MS and MS^2^) was then processed in Progenesis LC-MS v4.1 and the peptides identified in MASCOT database (Matrix Science,www.matrix-science.com).

### Statistical analysis

Statistical analysis was performed using the software Statgraphics Centurion ver. XV (Statpoint Technologies, Inc., Virginia, USA). Fitting of data was done using Origin ver. 8.1 (Origin Lab Corporation, MA, USA). Inhibition of biofilm formation data were fitted using a dose/response function (Boltzmann), according to the following equation (1):

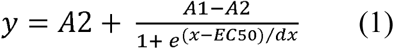

where A2 and A1 are the maximum and minimum level of biofilm formed, respectively, and EC_50_ is the concentration of the drug affecting 50% biofilm formation. EC_50_ values from the four replicates of the same drug were derived and their mean calculated. Distribution of EC_50_ values of the same drug was checked for normality. Analysis of variance (ANOVA) test was run to compare means from different drugs and to check variance. Probability value threshold was set to 0.05. Multiple comparison analysis was performed between pairs of data sets (considered as independent), corresponding to biofilm formation level (OD_490nm_) at each and every concentration of the given drug. F-test (ANOVA) was used to test for the rejection of the null hypothesis (P < 0.05).

## Results

### AHD drugs inhibited biofilm formation by S. mutans

AHD drugs did not show a visible growth inhibition effect in the concentration range studied (4– 2048 µg/mL). In the presence of sucrose, *S. mutans* sessile growth was not affected by any of the drugs, whereas all three AHD drugs inhibited biofilm formation in the presence of glucose, albeit with different potency (Fig. 1A). At concentrations >128 µg/mL of saxagliptin, there was 50% reduction of biofilm formation. Vildagliptin inhibited biofilm formation at concentrations ≥ 256µg/mL, while sitagliptin demonstrated at least 50% inhibition at 256µg/mL, albeit with a substantial increase in biofilm biomass at 2048µg/mL. The EC_50_ values of all drugs fell within the range of 128–512µg/mL. However, in the case of sitagliptin, the data point at 2048 µg/mL was not considered for fitting due to the anomalous behaviour at that concentration. While saxagliptin exhibited a more consistent inhibition pattern at concentrations ≥ 64 µg/mL, sitagliptin gave a sudden drop till saturation at 256–1024 µg/mL (Supplementary Fig.S1), as confirmed by the lower EC_50_ value (Table 1). In the case of saxagliptin and sitagliptin, the F-test (ANOVA) showed a statistically significant difference between the means at each concentration (p_saxagliptin_ = 0.0013, p_sitagliptin_ = 0.0089). To determine which means are significantly different from which other and to analyse the difference in effect at various concentrations, a multiple sample pairwise comparison of the means of the four independent measurements of biofilm formation at each drug concentration (number of concentrations n = 11, from 0 to 2,048µg/mL) was performed (Supplementary Fig. S2). The apparent increase in biofilm formation at 2048µg/mL of sitagliptin was found to be statistically relevant when compared with the biofilm biomass formed in presence of 128, 256, 512, and 1024 µg/mL of the drug (Supplementary Fig. S2b, cyan points). The differences observed in the presence of vildagliptin were not significant.

**Table 1.**
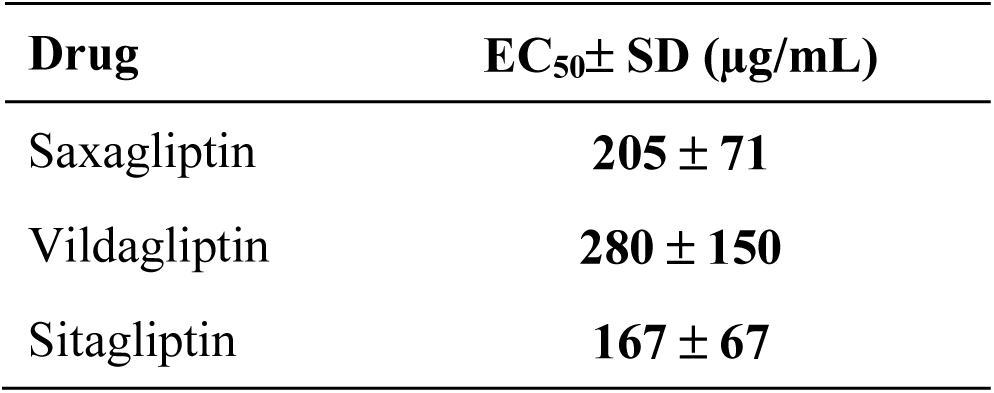
EC_50_ values for each drug. Mean concentration of drug producing 50% reduction in biofilm formation (EC_50_) was calculated from four independent replicates. Standard deviations (SD) were also determined.

**Fig. 1.**
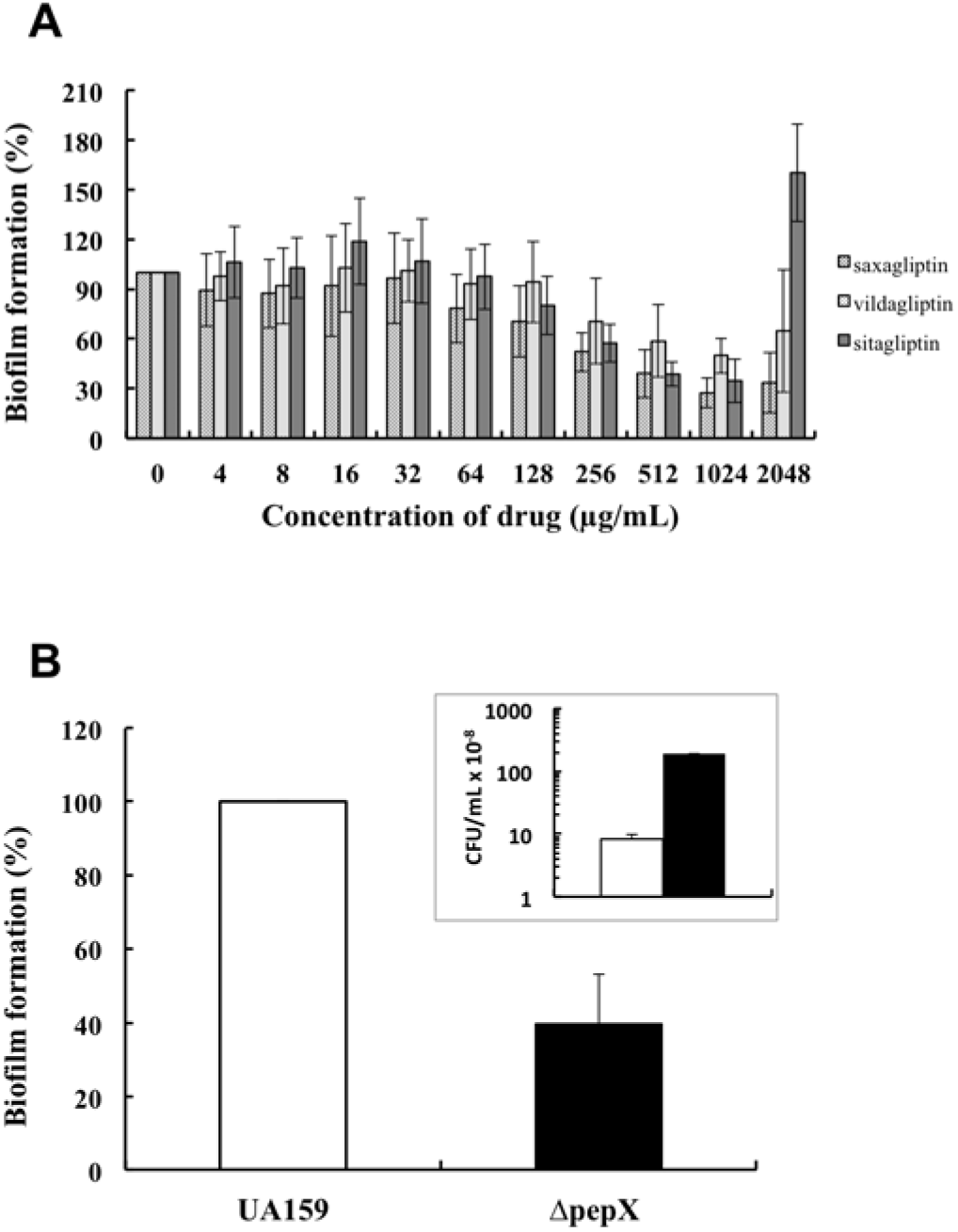
Quantification of biofilm formed by *S. mutans* in the presence of AHD drugs and of biofilm formed by a *pepX* deletion mutant. A. The histogram represents mean % biofilm formation in minimal medium containing 0–2048 µg/mL AHD drugs (n = 4, with duplicates within each independent experiment), B. Comparison of biofilm formation between Δ*pepX* (in black) and wild-type (in white) *S. mutans* in minimal medium containing glucose (n = 3, performed in triplicates within each independent experiment). Error bars represent standard deviation (p-value <0.001). Insert: viable cells count (expressed as CFU/mL) in the planktonic phase over the biofilm formed in SDM containing glucose.

The cells in the planktonic phase over the formed biofilm at each concentration of drug were equally viable with colony count at 64µg/mL-2048µg/mL of each AHD treatment ranging between 10^9^ to 10^10^ CFU/mL (Supplementary Fig. S3). The control group (8.23±1.7×10^8^CFU/mL) differed by one order of magnitude from the colony counts at 2048µg/mL of each drug (p = 0.043, unpaired student t-test, two-tailed).

### pepX mutant exhibited reduced biofilm biomass

The stronger effect of saxagliptin on biofilm compared to the other two AHD drugs, concomitant with a lower K_i_ of this drug against pure recombinant enzyme (De et al., 2016), suggested a probable role of *pepX* in influencing biofilm formation by *S.mutans*. To confirm a direct involvement of Sm-XPDAP in the biofilm development, a *pepX* mutant was generated. Crude extracts from the deletion mutant showed no amidolytic activity. The growth rate of the *pepX* deletion mutant was comparable to that of the wildtype both in BHI (data not shown) and inSDM(Supplementary Fig. S4, P = 0.317 by ANOVA for the variable “growth rate”), however, the CFU of mutant significantly differed from that of the wildtype hinting towards less viability of *pepX* mutant in SDM (Supplementary Fig. S4, P < 0.01 by ANOVA for the variable “growth rate”). The biofilm of the ∆*pepX* mutant in SDM containing glucose was reduced by 70% (SD = 13.4%, n = 9, Fig. 1B), while a lower decrease was observed in presence of sucrose (20%; SD = 3.8%, n = 3, data not shown). The planktonic phase over the biofilm of *pepX* mutant had more viable cells compared to wildtype (Fig 1B, insert), which may indicate that the mutant cells aggregate or adhere to the surface of the well less efficiently with a high propensity to remain in suspension. The same was for biofilms formed in the presence of AHD (Supplementary Fig. S3).

### Sm-XPDAP deficiency and saxagliptin treatment induced stress in S. mutans

The adherence of wild-type *S. mutans* treated with saxagliptin or vildagliptin and that of the Δ*pepX* mutant determined at 4h did not show any difference when compared to the untreated wild-type. At 20h, the wild-type demonstrated a thick and dense homogenous layer of aggregated cells, apparently more viable with few void spaces (15µm and 70.4% green cells, respectively), whereas the ∆*pepX* mutant exhibited a less organized thin layer of cells with more void spaces (9.6µm and 28.5% green cells)(Fig. 2). The effect of saxagliptin on wild-type was similar to that of ∆*pepX*. At 512 µg/mL saxagliptin, streptococci showed shorter dispersed chains and experienced apparent stress with a higher prevalence of propidium iodide stained cells (36%) (Fig. 3e and 3f). Vildagliptin also led to disaggregation of the biofilm compared to the untreated control, but streptococcal chains looked healthier (Fig.3g). *S. mutans* grown in the presence of 2048µg/mL sitagliptin presented a biomass level comparable to the untreated cells. However, chains were relatively long, apparently under stress with no difference in aggregation and in the proportion of dead cells (Fig. 3h &i).

**Fig. 2.**
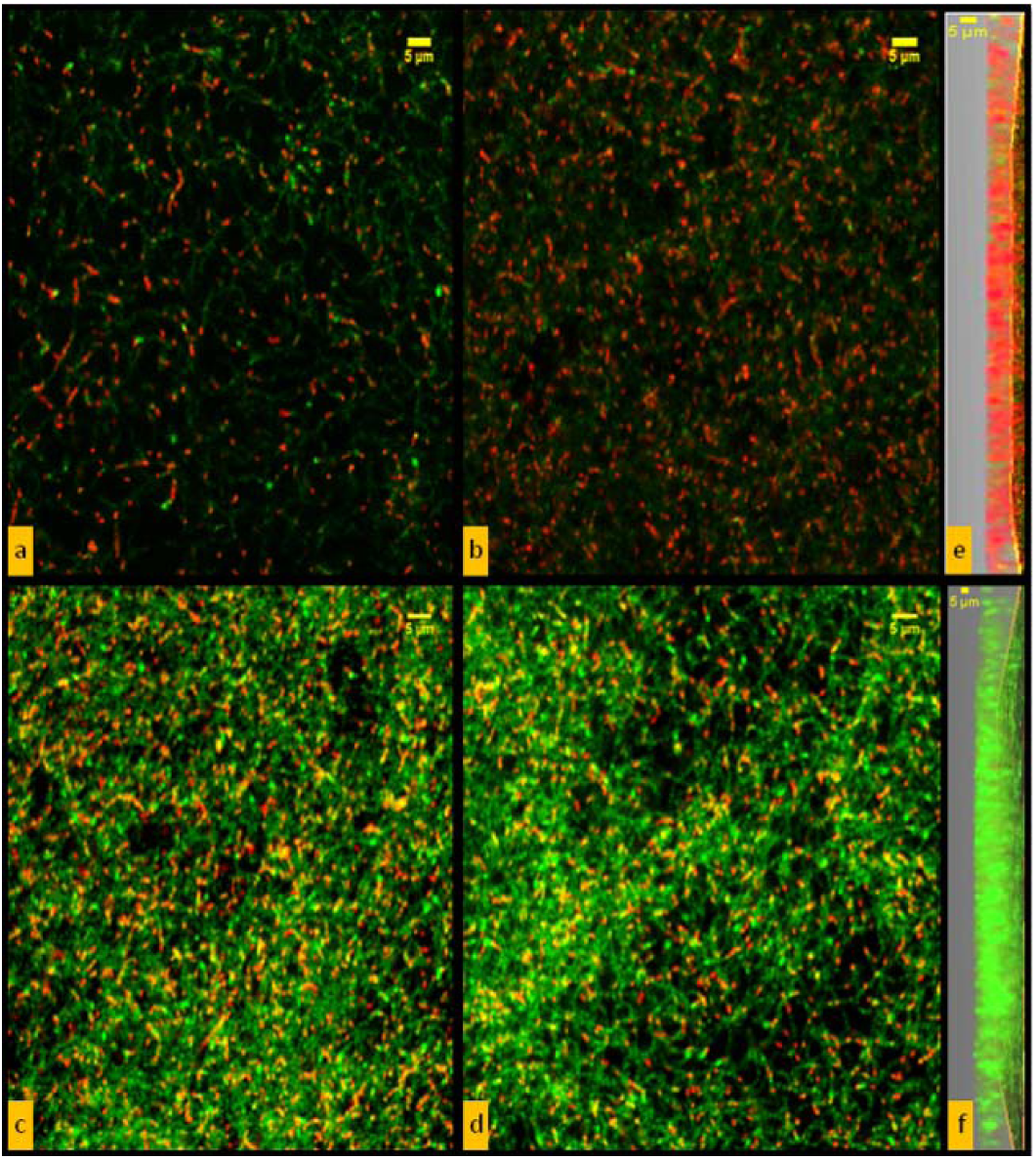
CLSM comparative study of biofilms formed by the ∆*pepX* mutant and wild-type *S. mutans* after 20 h. (a) ∆ *pepX* mutant (1^st^ field), (b) ∆ *pepX* mutant (2^nd^ field), (c) wildtype (1^st^ field), (d) wild-type (2^nd^ field), (e) z-axis representation of biofilm formed by ∆ *pepX* mutant, (f) z-axis representation of biofilm formed by wild-type.

**Fig. 3.**
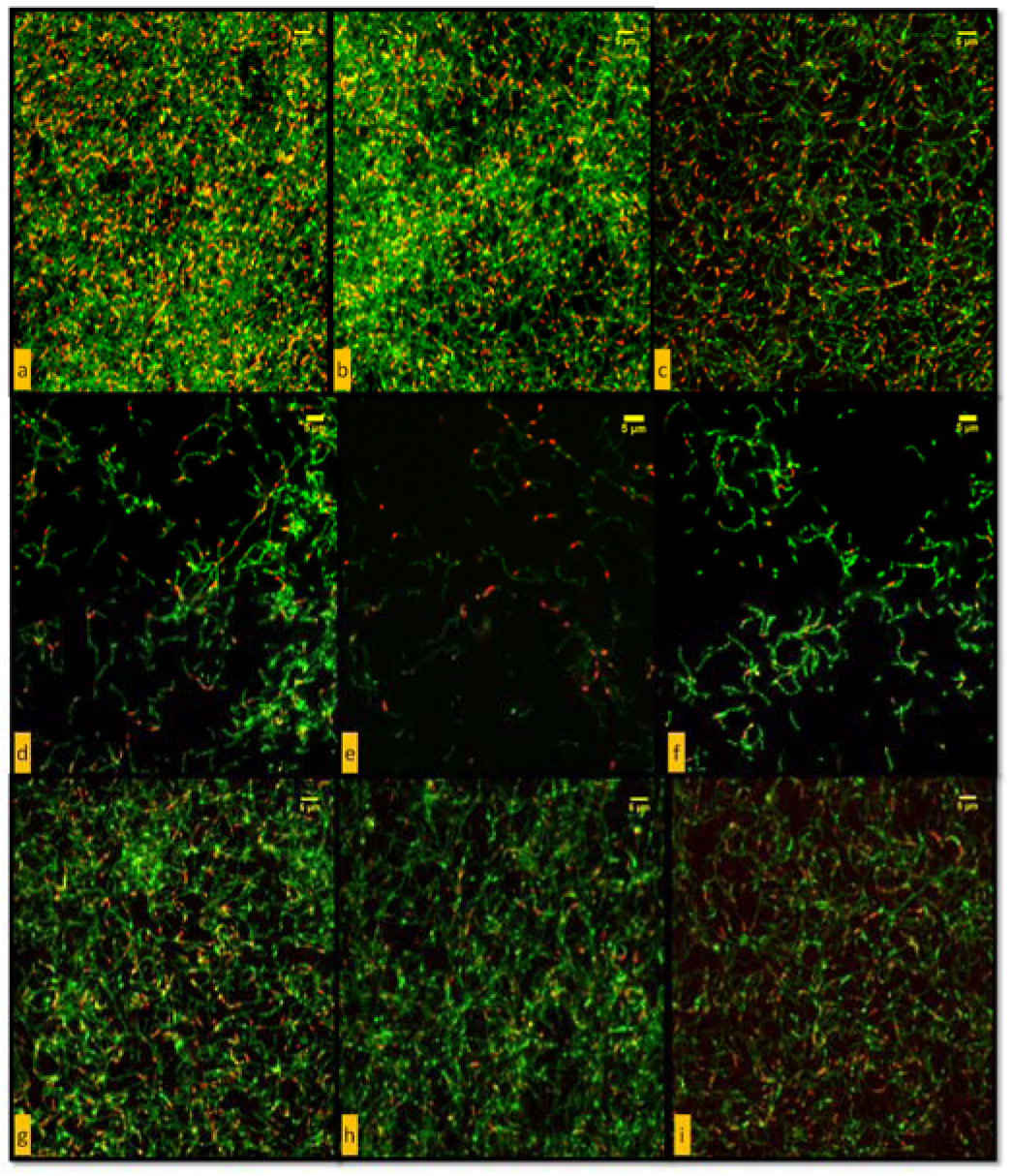
CLSM images of *S. mutans* untreated controls and treated with AHD drugs. (a) Untreated (1st field), (b) Untreated (2nd field), (c) Untreated (3rd field), (d) 128µg/mL saxagliptin treated, (e) 512µg/mL saxagliptin treated (1st field), (f) 512µg/mL saxagliptin treated (2nd field), (g) 512µg/mL vildagliptin treated, (h) 2048µg/mL sitagliptin treated (1st field), (i) 2048µg/mL sitagliptin treated (2nd field).

### pepX mutant exhibited downregulation of Cell Surface Antigen I/II

The proteomic analysis for differentially expressed proteins confirmed the absence of PepX in the Δ*pepX* mutant. Using relatively stringent criteria for protein identification (>1 peptide identified per protein in each biological replicate) only 9 proteins were found to be more abundant in the wild-type compared to the Δ*pepX* mutant (Supplementary Table S1) and no proteins were more abundant in the mutant compared to wild-type. Notably, the differentially expressed proteins included the Cell Surface Antigen I/II, a well-characterised adhesion protein of *S. mutans* (Jenkinson and Demuth, 1997). Notably, three of these 9 proteins were amino acid tRNA ligases. Comparison of control and sitagliptin-treated biofilms identified 6 proteins as more abundant in the controls and 26 as more abundant in the drug-treated cells. Likewise, comparison of control and saxagliptin-treated biofilms identified 6 proteins as more abundant in the controls (including three glycosyltransferases and levansucrase, which may affect biofilm matrix formation) and 23 as more abundant in the drug-treated cells. Two of the glycosyltransferases (SMU.910 and SMU.1005), upregulated in the controls compared to saxagliptin-treated biofilms, were upregulated in the sitagliptin-treated biofilms suggesting drug specific responses that may affect biofilm remodelling. Of 49proteins differentially expressed in either drug treatment, 12 were noted to be consistently altered in both (Supplementary Table S1) and were primarily involved in protein synthesis (n = 4) and various metabolic pathways (n = 7).

## Discussion

The AHD drugs used in this study are highly selective human DPP IV inhibitors (Wang et al., 2012). We hypothesize their effects on *S. mutans* by acting on Sm-XPDAP and regulating the behaviour of the bacterium, even though the drugs may have multiple enzyme targets. The *exvivo* assays using these gliptins showed inhibition of *S. mutans* biofilm formation. It could be speculated that this may impair streptococcal activity in the oral cavity of diabetic patients on these medications, as a consequence of the excretion of these drugs in saliva. Saxagliptin and vildagliptin have been found to possess 50% and >90% oral bioavailability, respectively (Boulton et al., 2013; Villhauer et al., 2003), while saxagliptin has also been reported in the salivary gland tissue (Fred, 2009). Sitagliptin, which is quite effective against biofilm formation and classified as a high intestinal permeability and low protein binding drug, can be detected in saliva (AUC 592 ng/mL×hr) (Idkaidek and Arafat, 2012). This may suggest the possibility of excretion of the other two AHD drugs in saliva. Furthermore, our results may suggest that systemic basal level of these drugs in host tissue of prescribed users may help in interfering with *S. mutans* behaviour.

In view of the differential cell physiology in sessile form compared to planktonic phase, the very high MIC of AHD drugs was considered inconsequential. CLSM was performed to investigate the influence of AHD drugs and the effect of Sm-XPDAP deficiency on the structural organization of *S. mutans* biofilms *in vitro*. We tested saxagliptin and vildagliptin, the first for its gradual relative effect at each concentration and the second for its consideration as a reference molecule for comparison between “cyanopyrrolidides”. CLSM was conducted at 128 µg/mL saxagliptin and 512 µg/mL vildagliptin, which were the concentrations causing >40% biofilm inhibition (Fig. 1A). Blurred green cells observed under CLSM in biofilm treated with AHD was indicative of the stress experienced by *S. mutans* in the given condition (Boulos et al., 1999). Cells treated with 512 µg/mL saxagliptin displayed similar observation to that of *pepX* mutant, which may allow hypothesizing that the inhibition of biofilm in presence of AHD drugs may be exerted through Sm-XPDAP inhibition. In contrast to the other AHD drugs, a sudden increase in attached biofilm observed in presence of 2048µg/mL sitagliptin can likely be best explained due to increased stress response. Interestingly, there was complete inhibition of growth at the subsequent higher concentration of sitagliptin (Supplementary Fig. S5). A similar differential response to varying concentrations of the antibiotic lincomycin has also been reported in *Streptococcus pyogenes* (Malke et al., 1981), albeit the mechanism of action has not been explained yet.

In this study a preliminary attempt was made to identify a plausible role of PepX in modulating the proteome and investigate the site of action of AHD drugs in *S. mutans* through whole cell proteome analysis. Considering the high amount of biofilm biomass required for proteome analysis in parallel to undergoing investigations of the observed effect of AHD molecules, the analyses were performed at 128 µg/mL of saxagliptin and sitagliptin. Vildagliptin was exempted from the study due to its limited effect as observed by CLSM, whereas use of 2048 µg/mL of sitagliptin was not feasible owing to the high amounts of pure molecule required. Surprisingly, relatively few differentially expressed proteins were consistently detected. Sitagliptin treatment and the *pepX* deletion both affected the expression of valine and proline amino acid tRNA ligases, which may reflect perturbation of cytoplasmic amino acid pools and consequently an overall stress as observed under CLSM. The alteration of the level of cell surface antigen I/II (Okahashi et al., 1989) was of much interest, and this may correlate with the reduced hydrophobicity and biofilm formation exhibited by the mutant. The 12 proteins that were differentially expressed in response to both drug treatments were noted to include enolase (SMU.1247), which can be a moonlighting surface-associated protein in *S. mutans*(Ge et al., 2004), and acetate kinase (SMU.1978), which participates in the Pta-Ack pathway that can influence biofilm formation.(Kim et al., 2015)

In conclusion, through this work we were able to establish a potential role of *pepX* in biofilm development of *S.mutans* and moved a step forward in the identification and development of novel Sm-DPP IV inhibitors. We have also tried to demonstrate the likely effect of AHD drugs on bacteria, as a proof of the concept that many regularly used drugs may exert some sort of side activity against microbes transiently or permanently inhabiting the human body. Future studies are focused on searching or synthesizing molecules targeting Sm-DPP IV that may be applicable as a new anti-caries agent.

## Acknowledgement

We acknowledge Prof. Giovanni Di Bonaventura and Prof. Simone Guarnieri at Università “G. d'Annunzio" Chieti-Pescara for their help in CLSM studies. This work was supported by a Grant from MIUR-ITALY - PRIN project 2009 no. 2009P5EKH4_003 (to G.L.) and by the University of Camerino grant no.FPA00057 (to V.L.A.).

## Conflicts of Interests

The authors declare no conflicts of interest.

